# Structural basis of multimodal adsorption and infection initiation by *Vibrio* phage Peru-2

**DOI:** 10.64898/2026.06.25.734500

**Authors:** Huaxin Yu, Justin Zhao, Jian Yue, Ian J. Molineux, Jun Liu, William P. Robins, John J. Mekalanos

## Abstract

Phage Peru-2, isolated during the 1993 cholera outbreaks in Peru, is distinct from the three ICP phage lineages typically associated with epidemic *Vibrio cholerae*. The molecular basis of Peru-2 adsorption and infection initiation has remained unknown. Here, we combine single-particle cryo-electron microscopy (cryo-EM) and cryo-electron tomography (cryo-ET) to define the architecture and infection mechanisms of Peru-2 at high resolution. The mature virion comprises an icosahedral capsid decorated with minor capsid proteins and a short tail apparatus surrounded by six structurally flexible tailspikes. These tailspikes are enzymatically active in mediating phage attachment to the *Vibrio* polysaccharide (VPS), a key component of biofilms. Three internal core proteins form a disordered core adjacent to the portal, positioning them for release before genome ejection during infection initiation. Structural analyses further resolve pre-ejection, genome-ejection, and post-ejection intermediates of the tail apparatus, while cryo-ET imaging of infected cells reveals a multimodal adsorption strategy during infection initiation.

**Significance:** *Vibrio* phages play important ecological and evolutionary roles, yet the structural basis underlying their host recognition and adsorption strategies has remained poorly understood. Here, we determine the overall architecture of *Vibrio* phage Peru-2 at near-atomic resolution, showing that it shares a conserved molecular organization with T7-like podophages but possesses additional minor capsid proteins and a distinct tailspike. In addition, Peru-2 interacts extensively with both the bacterial cell surface and sheathed flagella, revealing multiple modes of adsorption strategy distinct from those of classic T7 infection. Our structural analyses and functional evidence that *Vibrio* polysaccharide is required for Peru-2 adsorption and infection provide a mechanistic framework for understanding host recognition and infection strategies among *Vibrio* phages.

## Introduction

Cholera is a severely dehydrating diarrheal disease that has caused millions of deaths worldwide and remains a major global health threat, particularly in regions lacking access to clean water and adequate sanitation (1). The disease is caused by *Vibrio cholerae* (2), a highly motile Gram-negative bacterium that spreads primarily among humans through contaminated water. There are over 150 known serogroups, and only O1-based strains are responsible for the seven identified cholera pandemics (3). The current ongoing seventh pandemic, which began in 1961, is dominated by a group of related O1 El Tor strains (4).

*V. cholerae* motility is essential for navigation and bacterial fitness across diverse environments, including water and the host (5). During infection, motility enables *V. cholerae* to penetrate the intestinal mucus layer and reach epithelial niches (6), while nonmotile mutants exhibit a 100- to 1,000-fold reduction in colonization efficiency in the infant mouse model (7). *Vibrio* motility is driven by a single polar sheathed flagellum that is enclosed by a membranous sheath contiguous with the outer membrane and composed of lipopolysaccharide and outer membrane proteins (8, 9). In addition to motility, the sheathed flagellum contributes to surface attachment and initiation of biofilm formation in both the aquatic environment and small intestine during infection (10–12).

In contrast to its swimming and planktonic states, *V. cholerae* can transition to sessile, surface-attached growth within biofilms (13). Secreted *Vibrio* polysaccharide (VPS) is the major exopolysaccharide component of the biofilm matrix and associates with biofilm matrix proteins and extracellular DNA (eDNA) to form a three-dimensional protective barrier surrounding bacterial cells (14–16). Cells embedded within biofilms remain confined until *V. cholerae* expresses and secretes matrix-degrading enzymes that promote biofilm dispersal (17). Thus, *V. cholerae* has evolved to regulate both planktonic and sessile states, each of which contributes to bacterial fitness across diverse environments. As a protective barrier, biofilms shield *V. cholerae* from infection by commonly isolated ICP lytic bacteriophages (18). Moreover, *V. cholerae* can sense neighboring lysed cells and respond to phage-produced signals by enhancing biofilm production, representing a potential strategy to evade phage predation (19).

Lytic bacteriophages profoundly influence *V. cholerae* population dynamics in environmental reservoirs and during human infection (20, 21). In cholera-endemic regions, phage and bacterial counts in aquatic environments exhibit inverse seasonal correlations (20). Prominent lytic phages, such as ICP1, are amplified during outbreaks and reduce bacterial loads in environmental samples and patient stools (20, 21). Long-term monitoring of cholera outbreaks in Dhaka, Bangladesh identified three dominant phage lineages – ICP1, ICP2, and ICP3 – shed by patients (22). These phages effectively suppress *V. cholerae* colonization in infant rabbit and mouse models, particularly when administered as mixed cocktails (23, 24). Representatives of the ICP lineages have since been isolated from epidemics across multiple continents (25–29), underscoring the global relevance of phages as major drivers of *V. cholerae* evolution, ecological dynamics, and transmission.

Peru-2 is a *Vibrio* phage isolated during the 1993 cholera outbreaks in Peru. Although Peru-2 infects and lyses *V. cholerae* El Tor strains, it remains largely uncharacterized and is evolutionarily distinct from the well-studied ICP phages (30). Here, we report the annotated genome and structural characterization of Peru-2. Comparative genomic analyses reveal that Peru-2 shares a conserved genome organization and core genes with T7-like phages, including ICP3, while encoding additional features such as a second putative T7-like RNA polymerase and a unique tailspike protein. Using single-particle cryo-electron microscopy (cryo-EM), we determined near-atomic structures of Peru-2 in three distinct conformational states. These structures demonstrate that Peru-2 retains the canonical T7-like virion architecture while incorporating features reminiscent of *Salmonella* phage P22 (31), including a disordered internal core and six tailspikes containing a predicted carbohydrate-binding module (CBM) and a pectin lyase domain. Adsorption and infection are dependent on the production of VPS, which is likely recognized and degraded by the tailspike proteins. Complementary cryo-electron tomography (cryo-ET) of *V. cholerae* cells infected by Peru-2 reveals multimodal adsorption and infection intermediates *in situ*.

## Results

### The genome of Peru-2 resembles that of T7-like phages

The 46.6-kb genome of Peru-2 is organized similarly to, and shares extensive conserved gene content with, well-characterized T7-like phages (**Fig. 1A**). We selected T7 because it is closely related to ICP3, a group of phages commonly isolated during epidemics (22). Like other T7-like phages, Peru-2 encodes a single-subunit RNA polymerase (RNAP) expressed early during infection and predicted to direct transcription of essential viral genes (32, 33). Genes involved in DNA transcription, replication, virion assembly, structural morphogenesis, and genome packaging are arranged in modular clusters that closely mirror those of other T7-like phages, suggesting that the overall infection and intracellular replication processes are broadly conserved. Based on its genome size, organization, and gene content, Peru-2 is predicted to be an obligately lytic phage and shares numerous conserved functional and structural homologs with members of the order *Autographivirales*.

**Fig. 1.**
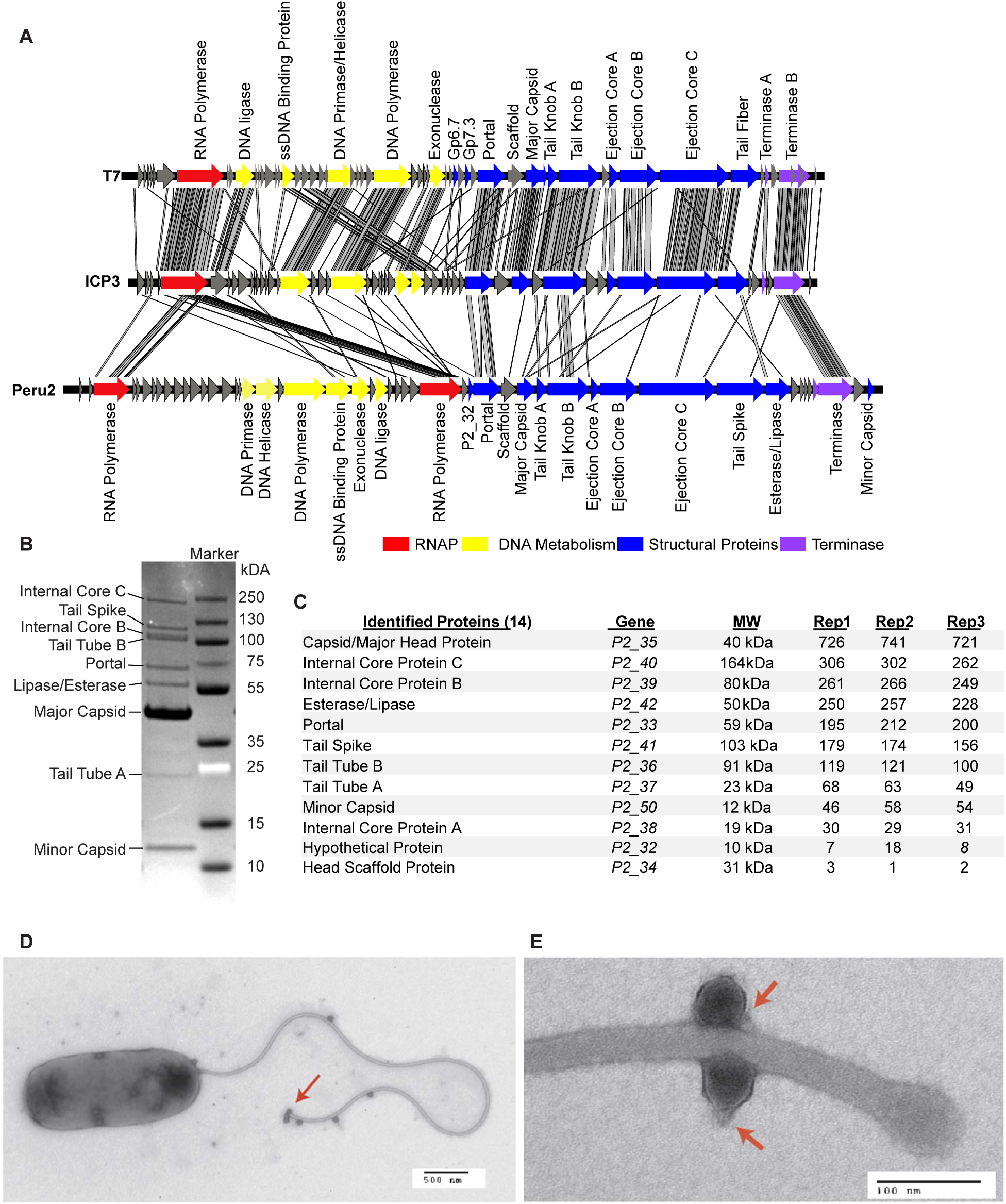
Characterization of *Vibrio* phage Peru-2. (**A)** Genome comparison of Peru-2 with T7-like ICP3 and T7. Gene protein products are colored by function. (**B**) This denaturing SDS-PAGE gel shows proteins from purified virions. **(C)** Peptides identified using mass spectrometry analysis from proteins extracted from purified virions. **(D)** Representative TEM images of *V. cholerae* cells infected by Peru-2. Numerous phage particles adsorb onto the sheathed flagellum and the cell surface. **(E)** Magnified TEM image showing two phage particles attached to the flagellum.

Despite this overall conservation, the early genomic region of Peru-2 shows notable divergence from T7 and *V. cholerae* T7-like phages such as ICP3. Most early genes in the first ∼25% of the genome lack clear homologs of canonical T7 early genes, several of which are nonessential or conditionally required for phage growth (34) (**Fig. 1A**). In addition, genes encoding orthologs of the two minor virion proteins gp6.7 and gp7.3 (35) are not readily identifiable in Peru-2 or ICP3, potentially due to their small size or the absence of recognizable sequence motifs. Nevertheless, several small open reading frames of unknown function with limited sequence similarity are positioned immediately upstream of the portal protein gene, suggesting analogous functions. In contrast to other T7-like phages, the modestly expanded genome size of Peru-2 is attributable in part to a gene encoding a second T7-like RNAP positioned between the DNA metabolism and structural modules (P2_30). Furthermore, in place of the canonical T7 tail fiber gene, Peru-2 encodes a substantially larger tailspike protein (P2_41) adjacent to a gene encoding a 473-amino-acid GDSL family lipase/esterase (P2_42).

To define the virion protein composition, purified Peru-2 particles were isolated by sequential CsCl step-gradient and equilibrium centrifugation and analyzed by SDS-PAGE and peptide mass spectrometry. SDS-PAGE revealed a limited number of prominent protein bands corresponding in apparent molecular weight to most predicted structural components, along with a highly abundant ∼12-kDa protein and an additional ∼55-kDa species (**Fig. 1B**). Peptide mass spectrometry of purified virions confirmed the presence of all predicted structural proteins as well as a high-abundance representation of lipase/esterase P2_42 and small protein P2_50 and low abundance of a protein encoded by gene *p2_32* (**Fig. 1C**). Notably, *p2_32* occupies a genomic position and size similar to T7 genes *6.7* and *7.3* (**Fig. 1A**) but is not readily visible on the stained gel, likely due to its small size and low abundance in the virion.

### Adsorption of Peru-2 to both the cell surface and flagellum

To determine the adsorption pattern of Peru-2 on host cells, we performed negative-stain electron microscopy of *V. cholerae* cells infected by Peru-2. Adsorbed phage particles were observed on the cell surface (**Fig. 1D**) as well as frequently along the polar sheathed flagellum (**Fig. 1E**), which is 1.5-4 times longer than the bacterial cell and can rotate at speeds up to ∼100,000 rpm (36). These observations indicate that Peru-2 likely recognizes a common receptor present on both the outer membrane and the sheathed flagellum.

### Cryo-EM structure of mature Peru-2 at near-atomic resolution

To define the architecture of Peru-2 and understand its adsorption and infection mechanisms, we used single-particle cryo-EM to analyze purified, vitrified Peru-2 virions. From 6,621 micrographs, 198,465 phage particles were selected to reconstruct a near-atomic structure of the mature virion (**Fig. 2A,B; *SI Appendix*, Fig. S1, S2**). The Peru-2 capsid is built from 415 copies of the major capsid protein P2_35, arranged into 60 hexamers and 11 pentameric vertices. P2_35 adopts the classic HK97 fold (37), closely resembling the T7 major capsid protein gp10 (38), with a root-mean-square deviation (RMSD) of 1.29 Å (***SI Appendix*, Fig. S3A**). In contrast to T7, the Peru-2 capsid is decorated at all 11 non-portal pentameric vertices by pentamers of the minor capsid protein P2_50 (**Fig. 2C; *SI Appendix*, Fig. S4A,B**). Each P2_50 subunit adopts an immunoglobulin (Ig)-like fold (***SI Appendix*, Fig. S4C,D**) consisting of seven β-strands arranged into two opposing β-sheets to form a β-sandwich followed by a C-terminal α helix, a motif frequently employed by tailed phages to mediate auxiliary interactions with host surfaces (39). The P2_50 pentamer forms extensive contacts with the underlying pentameric capsomer (**Fig. 2C,F**), suggesting roles in capsid vertex stabilization and potentially in host interaction, as observed for analogous Ig-like decorations in other phages (40).

**Fig. 2.**
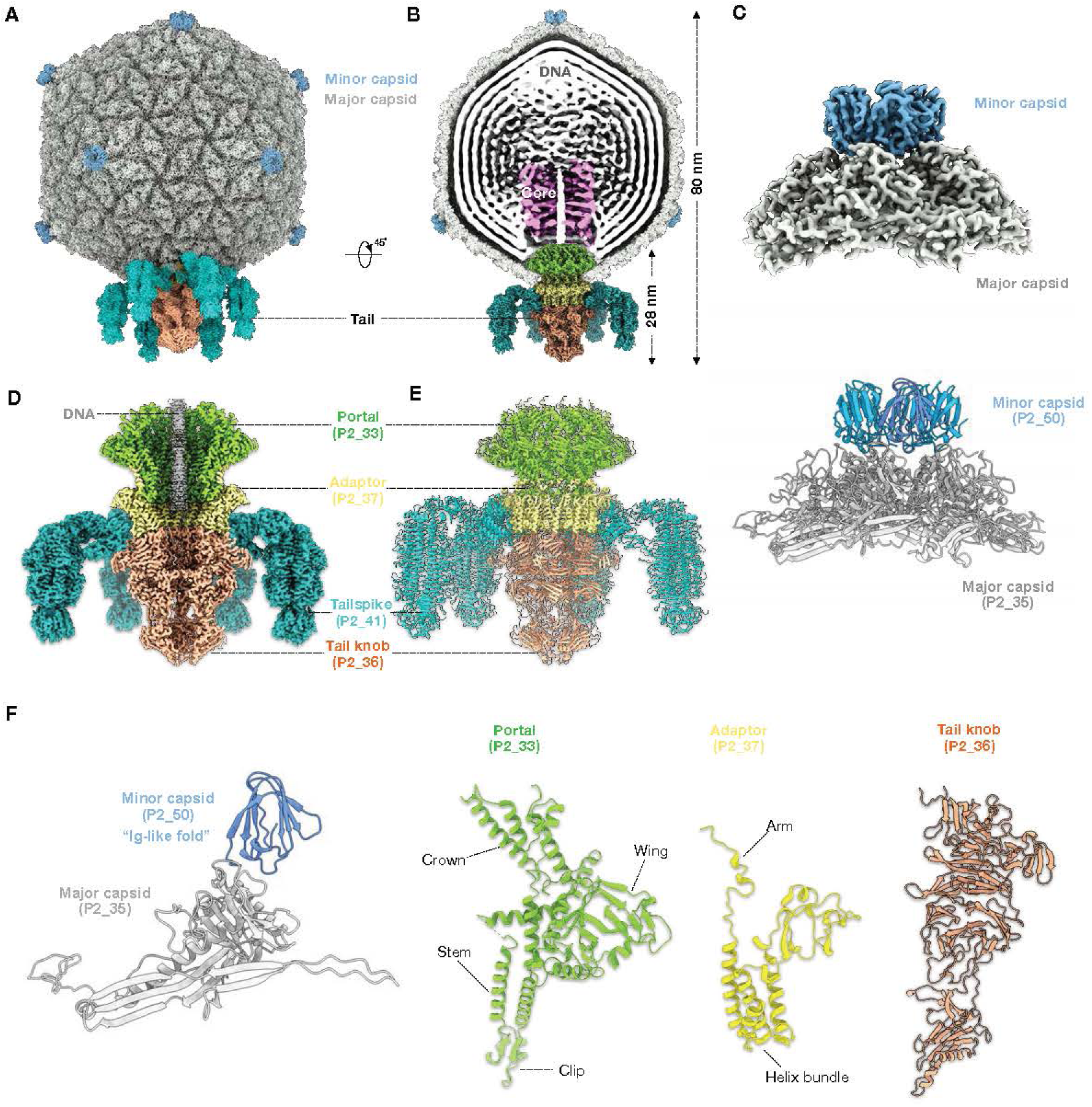
Overall structure of *Vibrio* phage Peru-2. **(A)** Tilted view of an intact Peru-2 virion. The icosahedral capsid features distinct minor capsid proteins decorating each of the 11 non-portal vertices. The remaining vertex is occupied by the tail apparatus, which is surrounded by six tailspikes. **(B)** Cutaway view of the complete Peru-2 virion, with the two frontal tailspikes omitted, revealing the protein shell that encloses the densely packed genomic DNA. A disordered internal core is visible directly above the portal-tail complex. **(C)** *Top:* Focused reconstruction of a pentameric capsid vertex in a side view, showing the minor capsid protein (P2_50) positioned apically on the major capsid protein (P2_35). *Bottom:* Ribbon representation of the P2_50 and P2_35 subunits adopting a fivefold symmetric arrangement. **(D)** Cutaway view of the focused reconstruction of the mature portal-tail complex, highlighting the closed tail knob and the extension of genome density into the portal-tail lumen, together with associated tailspikes. **(E)** Ribbon model of the complete portal-tail assembly. **(F)** Ribbon representations of individual subunits: Major capsid protein gp_35 interacts with minor capsid protein Gp_50, the portal protein P2_33 (residues 1-493, excluding the unresolved C-terminal portal barrel), the adaptor protein P2_37, and the tail knob protein P2_36.

At the unique portal vertex, a dodecameric portal complex composed of P2_33 forms the conduit for genome packaging and ejection (**Fig. 2D,E**). The portal protein adopts the canonical portal architecture (41), comprising crown, clip, stem, and wing domains, although the C-terminal domain of the portal (residues 494–530) is not resolved (**Fig. 2F**). Extending from the portal is a compact, non-contractile tail ∼28 nm in length. Twelve copies of the adaptor protein P2_37 assemble immediately beneath the portal, contacting it through extended arm helices, whereas six copies of P2_36 form the distal tail knob (**Fig. 2D–F)**. The tail lumen is markedly constricted and contains multiple closed gates (***SI Appendix*, Fig. S5A**). The narrowest constriction is formed by a loop spanning residues 504–510, where opposing Asp506 residues are separated by only ∼12 Å. In contrast to the portal and adaptor complexes, the tail knob exhibits substantial conformational flexibility, resulting in lower local resolution (***SI Appendix*, Fig. S1**). These dynamics are likely functionally important, as expansion of the tail knob is required to accommodate genome translocation during infection. Overall, the architecture of the portal-tail assembly closely resembles that of T7 (***SI Appendix*, Fig. S3B–D**), suggesting that Peru-2 and T7 share conserved mechanisms of virion assembly and infection initiation.

Peptide mass spectrometry identified three additional virion-associated proteins, P2_38, P2_39, and P2_40, that are homologous to the T7 internal core proteins gp14, gp15, and gp16, respectively. However, unlike the well-organized internal core formed by these proteins in T7 (35, 42), none of the corresponding Peru-2 internal core proteins is resolved. Instead, only weak density extending from the portal toward the center of the capsid is observed (**Fig. 2B**), suggesting that the Peru-2 internal core proteins occupy the defined region adjacent to the portal but remain largely disordered. In addition, the lipase identified as abundant by mass spectrometry is not observed in the cryo-EM structure, suggesting that it is either structurally disordered or degraded during phage maturation.

### Peru-2 encodes hybrid tailspikes that combine a T7-like tail fiber with a carbohydrate-binding and cleavage module

The tail of Peru-2 is surrounded by six trimeric tailspikes that form a prominent adsorption apparatus. In the consensus cryo-EM reconstruction of the tail complex, the tailspikes are only weakly resolved (***SI Appendix*, Fig. S1**), as previously shown for T7 tail fibers (43), indicating intrinsic flexibility. To resolve their structure, we performed focused classification and refinement, identifying a stable subset of particles that yielded a tailspike reconstruction at 3.65 Å resolution (***SI Appendix*, Fig. S1**), enabling *de novo* model building of the P2_41 trimer (**Fig. 3B; *SI Appendix*, Fig S2F**).

**Fig. 3.**
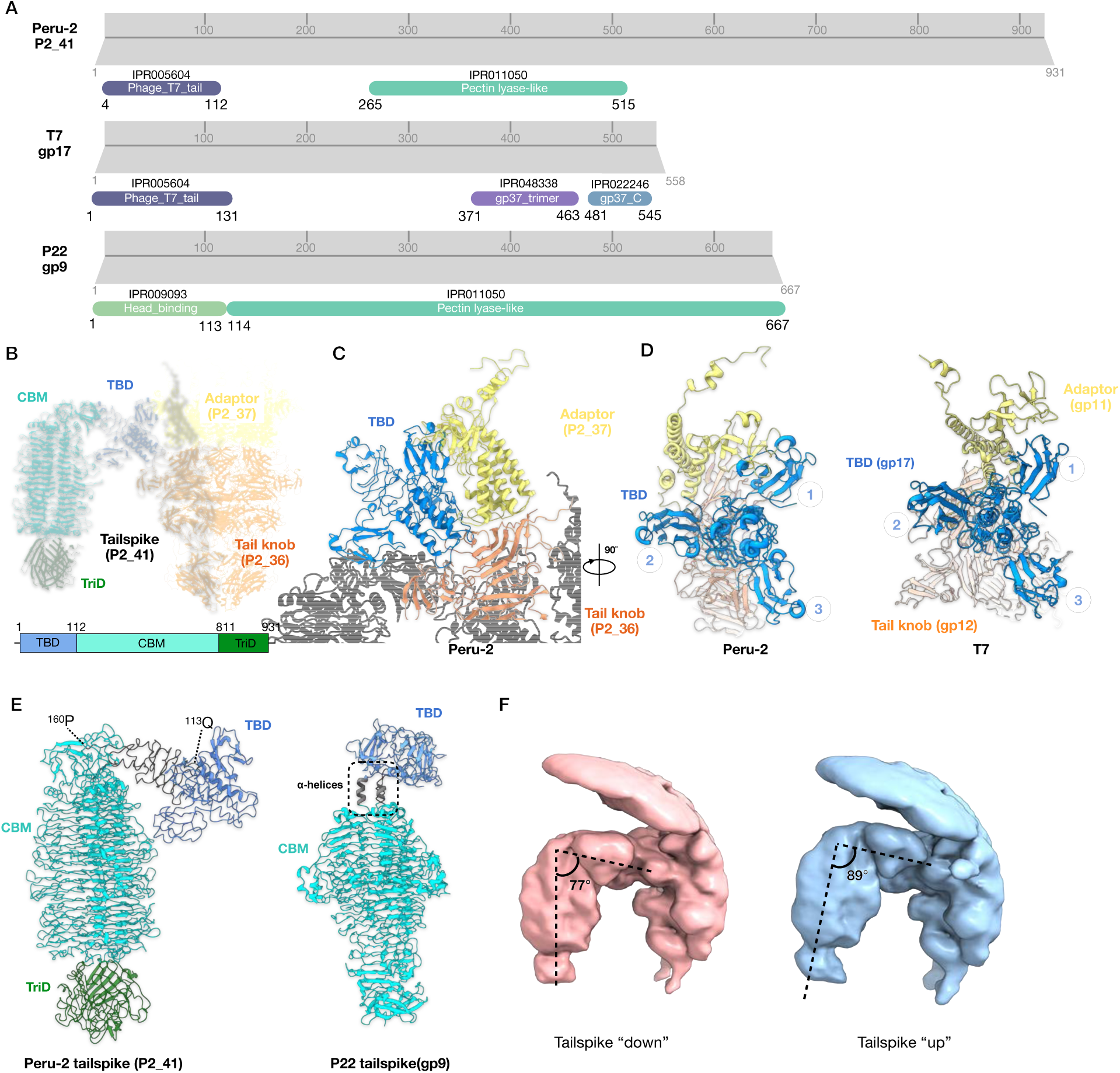
Architecture and dynamics of the Peru-2 tailspike. **(A)** InterPro domain annotations of Peru-2 P2_41, T7 gp17, and P22 gp9. **(B)** One Peru-2 tailspike shown together with its adjacent tail adaptor and tail knob. The tailspike is color-coded by domain: tail-binding domain (TBD, residues 1-112, blue) carbohydrate–binding module (CBM, residues 113-810, cyan), and trimerization domain (TriD; residues 811-931, green). **(C)** Left: Zoomed-in view of the joint region where the TBD interfaces with the tail adaptor and tail knob. **(D)** Central-axis views of the trimeric TBD from Peru-2 (left) and T7 (right, PDB: 9JYZ), each shown in complex with their corresponding tail adaptor and tail knob proteins. **(E)** Structural comparison of the Peru-2 tailspike internal domains, focusing on the parallel β-helix and triple β-prism. The Peru-2 structure is compared to phage P22 (PDB-8EAN). In Peru-2, the internal domains are connected by a flexible loop linker (residues 113-160), whereas P22 uses an α-helical connector at the analogous position. **(F)** Low-pass filtering of the Peru-2 tailspikes reveals two major conformations, termed tailspike “down” and “up”. The angles between the tailspike shaft and the head-binding domain measure 77° and 89°, respectively.

Each P2_41 monomer is organized into three distinct regions: an N-terminal tail-binding domain (TBD, residues 1–160), a carbohydrate-binding module (CBM, residues 161–810), and a C-terminal domain (TriD, residues 811–931). InterPro analysis identified a T7 tail fiber protein-like, N-terminal domain (IPR005604) spanning residues 4 and 112 and a pectin lyase fold/virulence factor domain (IPR011050) spanning residues 265 and 515 (**Fig. 3A**). The N-terminal tail-binding domain inserts into the symmetry-mismatched interface formed by the tail adaptor and tail knob complexes (**Fig. 3B**). Structural comparison further reveals that this domain closely resembles the tail-binding domain of the T7 tail fiber gp17 (42) (**Fig. 3C,D**), supporting a shared evolutionary origin.

The predicted CBM and pectin lyase domain of P2_41 are embedded within a long 18-turn parallel β-helix region (residues 161–810), a hallmark of other enzymatic tailspike proteins in many phages (44). Tailspike-associated polysaccharide depolymerases typically belong to either hydrolase or lyase families, which are specialized to recognize and degrade distinct polysaccharide substrates and glycosidic linkages present in bacterial antigens, capsules, and extracellular exopolysaccharides (45). In contrast to hydrolases, polysaccharide lyases cleave α- and β-1,4-glycosidic bonds through a β-elimination mechanism (46).

Because the *V. cholerae* O1 antigen is a homopolysaccharide composed of α-1-2 linked 4-amino-4,6-dideoxy-D-mannose (perosaminyl) residues (47, 48), we reasoned this was an unlikely receptor and substrate for the putative P2_41 tailspike pectin lyase domain. Instead, we considered VPS, the primary extracellular matrix component of *V. cholerae* biofilms, due to its molecular composition and linkages. A primary component of VPS is a tetrasaccharide repeating unit containing glucose, galactose, N-acetylglucosamine, and mannose with additional N- and O-acetylations and glycine linkages (***SI Appendix*, Fig. S6H)** (49). The α-1,4 linkages in VPS are cleaved by the endogenous *V. cholerae* lyase RbmB, which also features a parallel β-helical fold typical of the pectin lyase family, when the bacteria degrade VPS-biofilms to allow bacterial release from a sessile to planktonic state (50). Other phages degrade extracellular polymeric substances (EPS) in bacterial biofilms using tailspikes that possess pectin lyases (45).

To investigate potential VPS recognition, we employed Protenix, an open source reproduction of AlphaFold that accepts Protein Data Bank (PDB) molecules (51) to model the interaction of VPS and the tailspike trimer (***SI Appendix*, Fig. S6A–E,G,H)**. In all modeling iterations, the VPS ligand occupied this same binding site (***SI Appendix*, Fig. S6A–E**). In addition to 22 residues predicted to form H-bonds with this VPS and several aromatic residues, we identify two Lysine and three Arginine residues predicted to form salt bridges. These features are consistent with catalytic and substrate binding residues observed in other pectin lyase domains, which mediate substrate binding, stabilization, and cleavage during β-elimination reactions (52, 53). Independently, we used GlycanInsight, a deep learning-based platform that predicts carbohydrate-binding residues and pockets on protein structures using defined glycan ligands (54). Using the minimal VPS tetrasaccharide unit as the ligand, GlycanInsight similarly identified residues lining the outfacing surface in the CBM pectin lyase fold/virulence factor domain. Many of these residues overlap those predicted by the Protenix models (***SI Appendix*, Fig. S6F)**, collectively supporting a role for this domain in VPS recognition and catalysis. At its distal end, P2_41 forms a triple β-prism architecture, with each prism composed of an eight-stranded β-sandwich (**Fig. 3E; *SI Appendix*, Fig. S2F,G**). This distal β-prism may further engage inner- or outer-core lipopolysaccharide components or membrane proteins, thereby stabilizing irreversible phage attachment after initial VPS recognition.

Consistent with this proposed role in host engagement, the Peru-2 tailspikes exhibit pronounced flexibility. Focused classification resolves two predominant conformations **–** “tailspike-down” and “tailspike-up” **–** that differ by ∼12° at the junction between the tail-binding domain and O-antigen-binding domain (**Fig. 3F; *SI Appendix*, Fig. S1**). This flexibility is mediated by a short hinge region (residues 113–160) and likely increases the effective capture radius of the phage, thereby enhancing the probability of productive host contact. Together, these results show that the Peru-2 tailspike is a modular, multifunctional adsorption apparatus that combines T7-like attachment modules with carbohydrate-binding and catalytic activities.

### VPS is recognized by Peru-2 and required for infection

We first plated Peru-2 on *V. cholerae* isogenic strains with well-characterized Inaba, Ogawa, and Hikojima variants in the O1 antigen, as well as on mutants lacking flagella (55, 56). We observed little difference in plating efficiency relative to wild type (not shown). To test the role of VPS in phage adsorption and infection, we compared a mutant defective in VPS production with a mutant that constitutively overexpresses VPS, even during planktonic growth outside of biofilms. Peru-2 plated on a lawn of the VPS-defective mutant exhibited a ∼50-fold reduction in titer and produced only turbid plaques (**Fig. 4A)**. In adsorption assays, phage particles failed to adsorb to the VPS-defective mutant and adsorbed nearly 5-fold better than C6706 on an isogenic mutant that constitutively produced VPS after 30 minutes (**Fig. 4B,C**). Although TEM showed phage particles associated with cells and flagella, these results indicate that VPS production is required for irreversible adsorption and efficient Peru-2 infection.

**Fig. 4.**
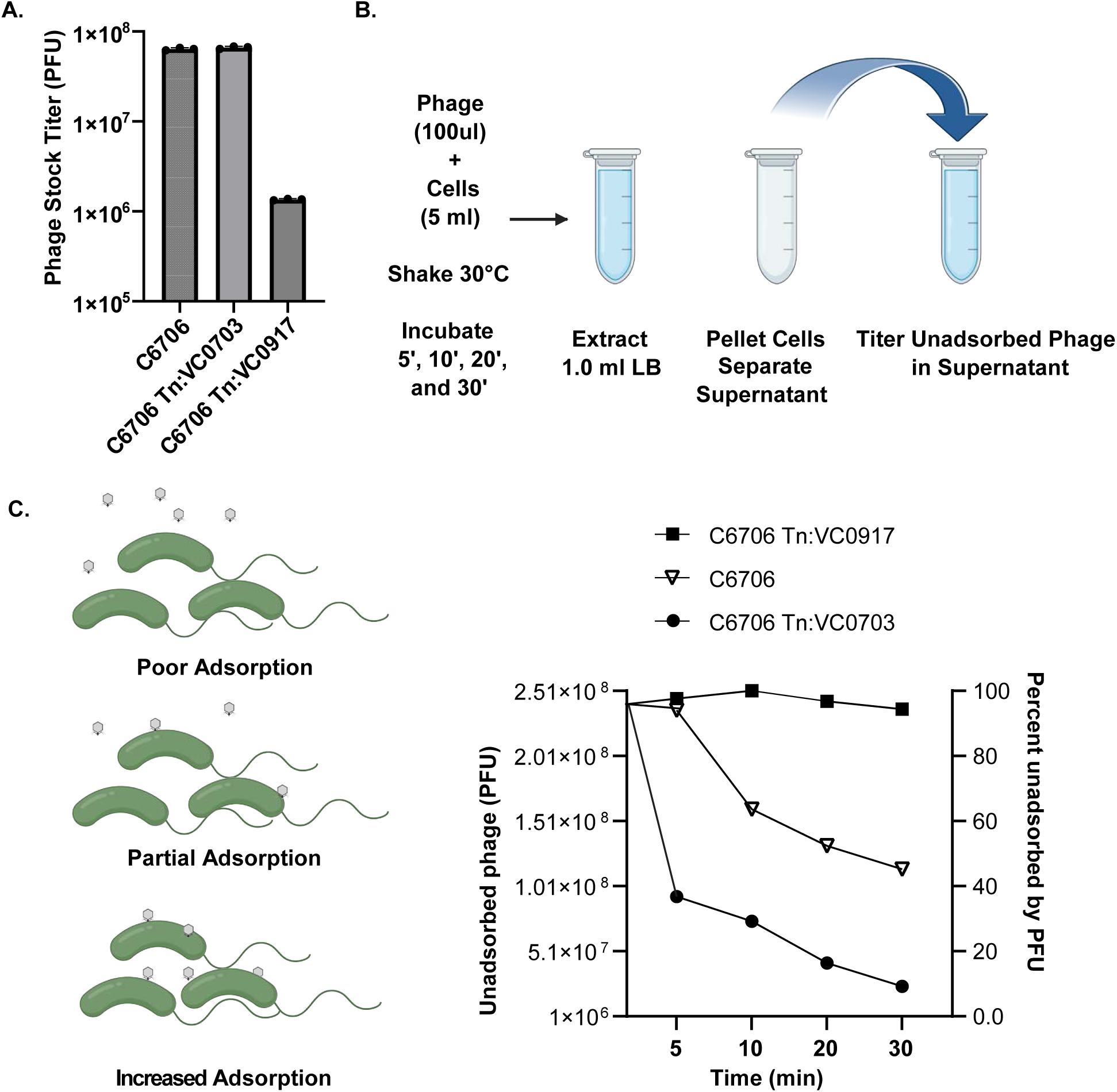
Phage bacterial adsorption and infection is largely dependent on VPS. (**A**) Peru-2 phage stock titered on C6706 and isogenic mutants that either constitutively or fail to produce VPS. (**B**) Experimental design of adsorption assay. (**C**) Model of phage adsorption and phage adsorption to C6706 and isogenic mutants that either constitutively or fail to produce VPS.

### Conformational changes and extension of the tail complex

Following adsorption, Peru-2 must undergo major structural rearrangements to enable the ejection of internal core proteins and then subsequent release of the genome. In addition to the closed tail knob observed in the mature virion, focused classification and refinement revealed a distinct open-tail conformation at 3.86 Å resolution, characterized by a pronounced expansion of the tail knob with a lumen diameter of ∼32 Å compared with ∼12 Å in the mature particle. In this state, the portal and adaptor remain unchanged (**Fig. 5A–C**), and the genome end appears confined within the portal-tail lumen (**Fig. 5D**), suggesting that this conformation represents an infection intermediate after translocation of internal core proteins but before genome release. This expansion results from an ∼7° outward rotation of each tail knob subunit relative to the virion axis (**Fig. 5E**). The arrangement is accompanied by the appearance of an additional density corresponding to P2_38 ejected from the capsid. Structural modeling of P2_38 residues 33–133 reveals a helix–turn–helix fold, with two long helices (residues 33–79 and 80–133) closely matching the conformation of the ejected gp14 in T7 (42). Notably, the N- and C-terminal regions of P2_38 remain unresolved but are predicted, based on the existing T7 model (42), to insert into the host membrane.

**Fig. 5.**
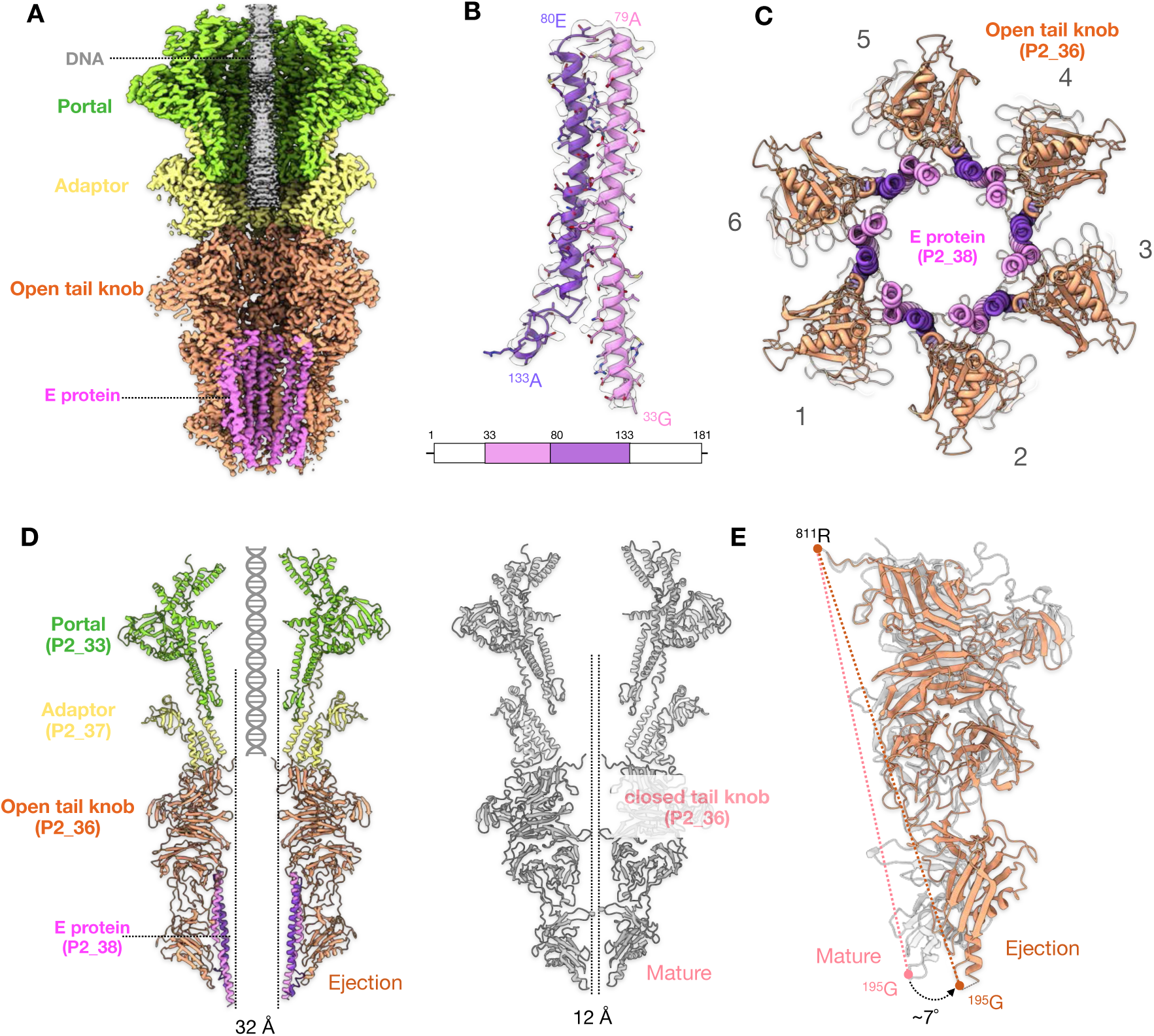
Tail opening and extension through interaction with internal core protein P2_38. **(A)** Cut-through view of the portal-tail complex from Peru-2 particles displaying tail open and extension. The packaged DNA remains inside the capsid, while an internal core protein P2_38 is ejected from the capsid and is partially visible in the lumen of the opened tail knob. **(B)** The P2_38 monomer adopts an intra-chain coiled-coil structure composed of two α-helices, with residues ^33^G–^133^A resolved in the density. **(C)** Bottom view of the opened tail knob in cartoon representation. **(D)** Side views comparing the portal-tail assemblies in the open (left) and closed (right) conformations. The open state exhibits a markedly widened knob aperture, with the tail knob P2_36 density expanding from ∼12 Å to ∼32 Å. **(E)** Superposition of the P2_36 monomer from both states reveals an approximately 7° rotation of the tail knob tip at residue Arg195 during transition from the closed to the open conformation.

Although our data do not establish whether tail knob expansion triggers P2_38 engagement or vice versa, their co-occurrence supports a model in which Peru-2 undergoes coordinated rearrangements analogous to the early steps of T7 infection initiation. These conformational changes transform the closed tail knob into an open channel sufficiently wide to accommodate and facilitate DNA passage (**Fig. 5D; *SI Appendix*, Fig. S6A,B**).

### Observation of the portal barrel after DNA ejection

We further resolved a third conformation of Peru-2 at 3.92 Å resolution, in which the viral genome has been fully ejected from the capsid while the tail knob remains in the open configuration (**Fig. 6A,B; *SI Appendix*, Fig. S2**), likely representing a post-ejection state. Notably, a well-defined portal barrel formed by an α-helix spanning residues 494–514 of P2_33 was observed (**Fig. 6C**). This barrel constricts the central pore of the portal from ∼58 Å to ∼39 Å in diameter (**Fig. 6D**). By contrast, this barrel density was not observed in the two pre-ejection conformations, suggesting that the portal barrel is either highly flexible or obscured by the packaged genome and internal core proteins. Following ejection of the internal core proteins and release of the genome, the portal barrel adopts a stable closed configuration that appears to seal the portal after DNA translocation.

**Fig. 6.**
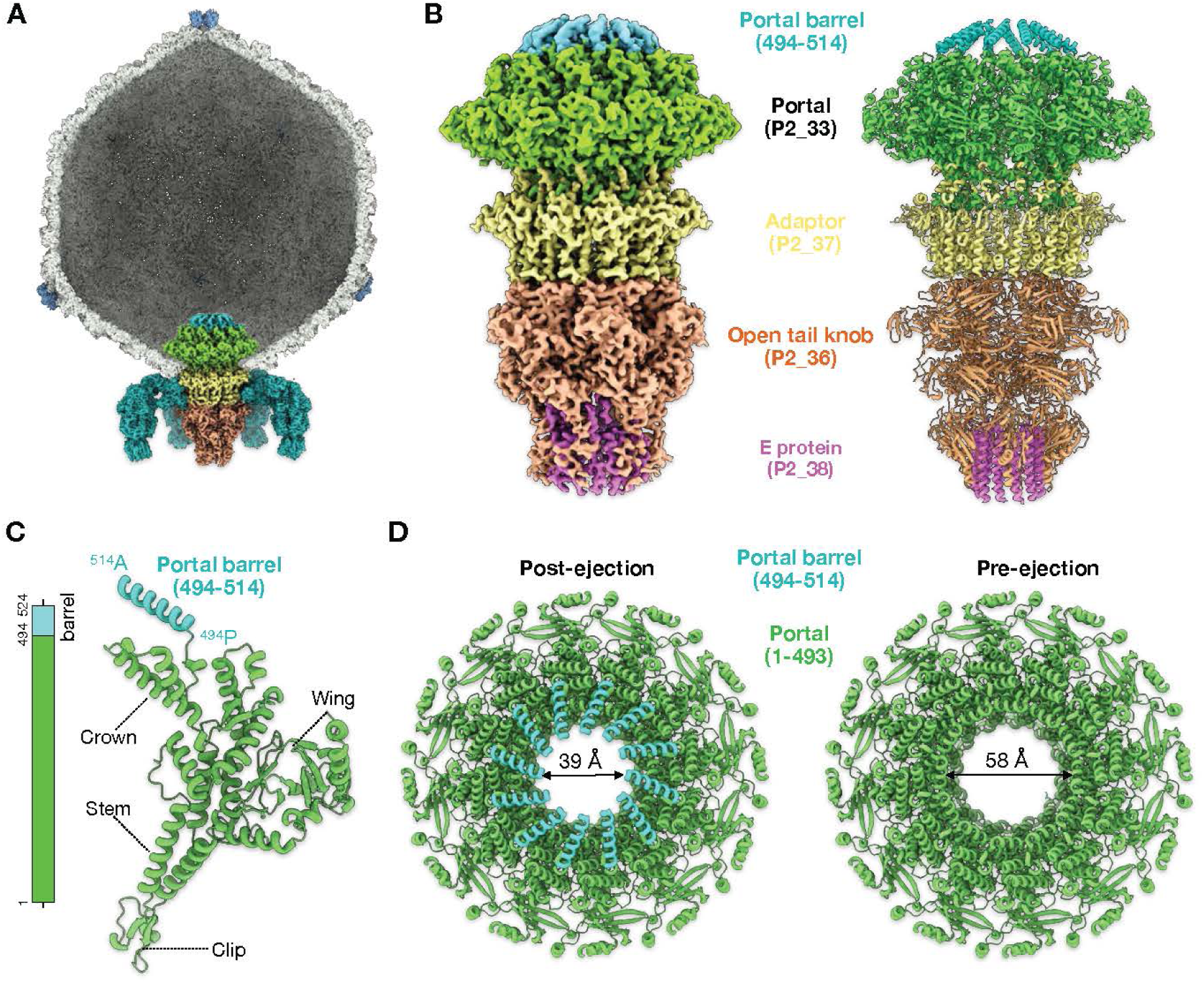
Peru-2 architecture after genome ejection. **(A)** Cut-through view of a Peru-2 virion with an empty capsid and the associated portal-tail complex; the two front-facing tailspikes are omitted for clarity. **(B)** *Left:* Local reconstruction of the portal-tail assembly in its post-ejection conformation. *Right:* Corresponding atomic model, highlighting the reformation of the portal barrel by P2_33 (cyan). **(C)** Structure of an individual P2_33 subunit, showing the α-helical barrel segment (residues 494-514) that becomes ordered in the post-ejection state. **(D)** Top views of the P2_33 portal complex in the post- and pre-ejection states, illustrating how the 12-fold symmetric portal barrel constricts to nearly seal the portal-tail lumen after genome release.

### Visualizing Peru-2 infection initiation by cryo-ET

Our single-particle cryo-EM analyses identified three distinct conformations of Peru-2 that likely correspond to pre-ejection, genome-ejection, and post-ejection states. To relate these structural states to infection initiation *in vivo*, we performed cryo-ET analyses on *V. cholerae* wild-type and Δ*flhG* mutant cells infected with Peru-2 (**Fig. 7; *SI Appendix*, Fig. S8A).** As expected, many adsorbed virions attach directly to the cell surface, presumably through interactions between tailspikes and outer membrane components (**Fig. 7C**). Upon engagement of an appropriate surface receptor, Peru-2 is expected to eject its internal proteins from the capsid, driving tail knob expansion and assembly of a transenvelope channel, analogous to the extended tail formed during T7 infection (43). Following genome translocation, the portal undergoes a conformational rearrangement to form a stable barrel, presumably sealing the conduit and preventing leakage of host cytoplasmic contents.

**Fig. 7.**
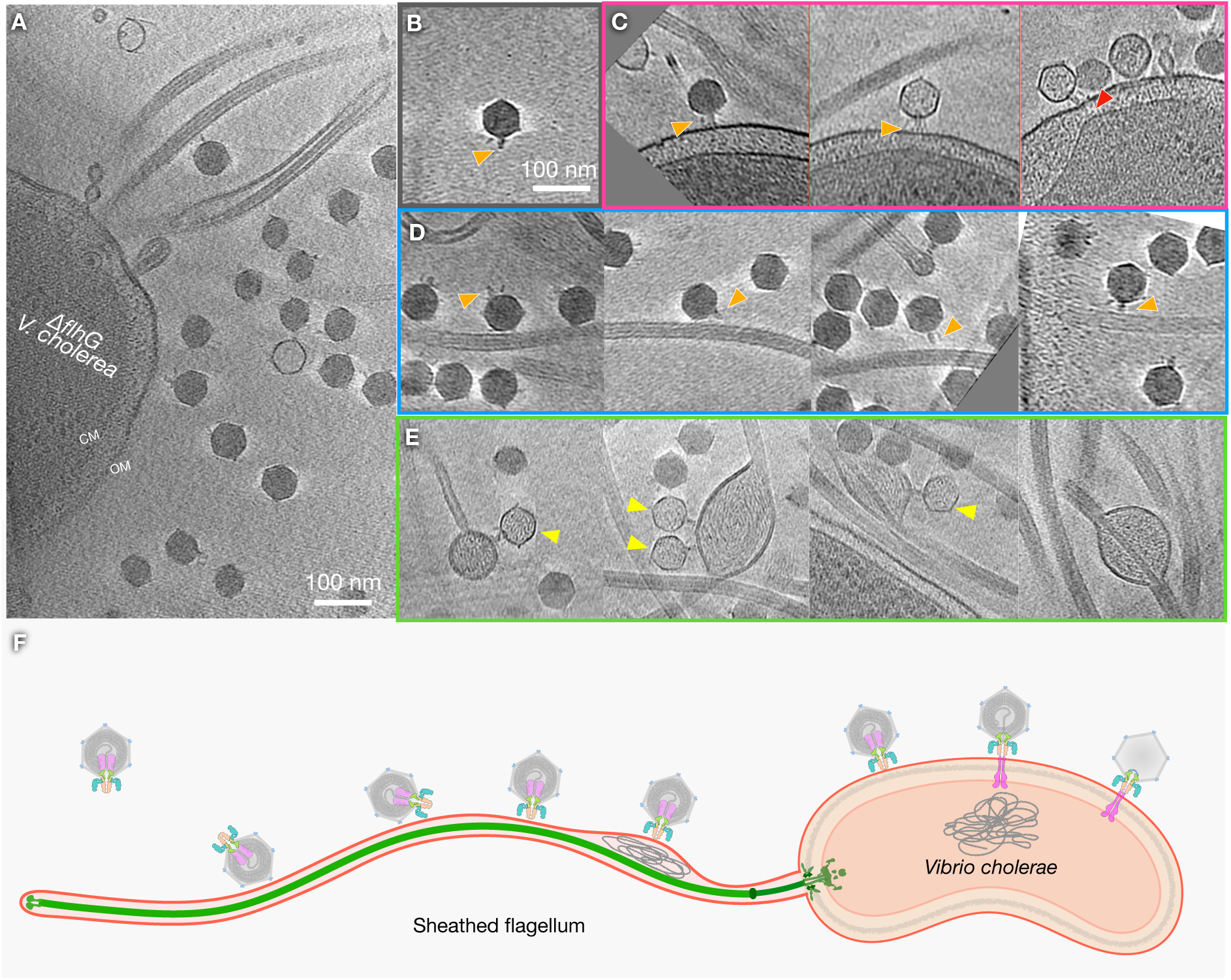
Overall schematic of Peru-2 infection initiation. **(A)** Representative tomographic slices of infection intermediates. **(B)** Free phage particle**. (C)** Sequential phage–cell membrane interactions: initial attachment (left) followed by genome release (middle and right). **(D)** Phage interacting with the flagellum via head and tail. **(E)** Peru-2 particles ejecting their genome into the sheathed flagellum or into vesicles associated with the flagellar sheath. Orange arrow, phage tail; red, trans-envelope channel. **(F)** Cartoon schematics illustrating the inferred infection state.

Cryo-ET also revealed extensive adsorption of Peru-2 onto sheathed flagella (**Fig. 7D; *SI Appendix*, Fig. S8B**). In the Δ*flhG* mutant, which produces increased numbers of aberrant flagella, both sheathed and unsheathed (57), the abundance of flagella-bound virions was markedly high (**Fig. 7D; *SI Appendix*, Fig. S8B,C**). Many particles appear to interact with the sheathed flagella via minor capsid proteins (**Fig. 7D; *SI Appendix*, Fig. S8B,C**), consistent with the model in which Ig-like modules among some tailed phages mediate auxiliary host-surface contacts (39). These multivalent, low-affinity binding interactions likely enhance encounter frequency in complex aquatic environments. Given that *V. cholerae* sheathed flagella rotate at high speeds (36), they may function as “phage collectors” promoting phage adsorption.

Notably, empty capsids were detected not only on the bacterial surface but also associated with sheathed flagella (**Fig. 7C,E**). These observations suggest that, under certain conditions, sheathed flagella alone may be sufficient to trigger genome ejection, revealing a previously unrecognized flagellotropic interaction mediated by the flagellar sheath of *V. cholerae*. However, it remains unclear whether genome release into the flagellar sheath can lead to productive infection. Taken together, these observations support a model of Peru-2 infection initiation in which adsorption to the cell surface or sheathed flagellum is followed by particle reorientation, tail extension, and genome delivery (**Fig. 7F**).

## Discussion

Peru-2 and T7 share conserved architectural features, reflecting a common evolutionary origin among T7-like podophages, including the ICP3 phages. These phages are built around an icosahedral capsid containing a linear dsDNA genome, a dodecameric portal at a unique vertex, internal core proteins, and a short tail apparatus that mediates genome translocation across the bacterial envelope. These conserved elements define a common infection strategy in which internal core proteins are ejected first to assemble a transenvelope conduit, followed by release of the genome into the host cytoplasm. Our structural analyses confirm that Peru-2 shares this model, including the internal core and short tail apparatus, both essential for genome delivery.

Peru-2 diverges from T7 in how it engages and penetrates the host cell surface. Whereas T7 employs flexible tail fibers for reversible adsorption to surfaces and increased receptor engagement in cells (58), Peru-2 instead encodes prominent tailspikes. Structural homology and experimental evidence strongly suggest that Peru-2 tailspikes recognize and enzymatically degrade VPS in biofilms, thereby clearing a path to the cell surface. Infection is partially VPS-dependent, as phage particles adsorb poorly to planktonic cells growing in liquid media unless the strain constitutively produces VPS.

Other characterized phages that also utilize tailspike depolymerases to degrade capsular and other extracellular polysaccharide antigens likewise often require these structures for efficient infection (59). Consistent with this idea, we previously observed that liquid culture-grown bacterial cultures are never completely lysed by Peru-2, even at high MOI, whereas plate-grown bacteria embedded at high density in soft agar support efficient phage propagation. The laboratory strain C6706 used to propagate Peru-2 carries a *luxO G333S* allele that partially enhances biofilm formation through constitutive low-level VPS production (60). Similar *luxO* mutants arise independently during laboratory passage of *V. cholerae* strains, including C6706, and appear to increase both bacterial fitness under laboratory conditions and Peru-2 adsorption efficiency growth (60). In newly isolated clinical strains prior to laboratory passage, *luxO* mutants are absent. We therefore reason a stringent regulation of VPS production may normally protect planktonic cells from infection by phages such as Peru-2. Conversely, biofilm-dependent phages may be particularly advantageous in environments where biofilm growth is induced.

A second distinction between Peru-2 and T7-like phages is the presence of the minor capsid protein P2_50, which contains an Ig-like fold decorating the non-portal vertices of the Peru-2 capsid. Some T7-related phages synthesize a small fraction of major capsid protein variants altered in the C-terminus via frameshift, and these minor capsid proteins introduce additional amino acids at some vertices (61, 62). In T3, the extended minor capsid protein contains a predicted Ig-like domain that interacts with mucins (39, 63) although no direct enhancement of bacterial adsorption has been demonstrated. Accordingly, the Peru-2 minor capsid protein may serve dual roles in stabilizing the capsid and in maintaining virion proximity to bacterial cells. This interpretation is consistent with the widespread distribution of Ig-like domains among tailed phages, where they frequently function as accessory adsorption modules under complex environmental conditions, including mucosal environments where bacteria and phages co-localize (63).

We further propose that phages reversibly adhering to planktonic cells or flagella may subsequently be transported to surface-attached cells residing within biofilms. The *V. cholerae* sheathed flagellum plays a central role in biofilm development by mediating the initial attachment of cells to abiotic surfaces (10). Furthermore, the flagellum acts as a mechanosensor capable of stimulating VPS production through cyclic-di-GMP signaling when flagellar function is impaired (12, 64). Although we cannot determine the precise organization or abundance of VPS associated with planktonic cells and flagella, we reason that reversibly attached phages may gain access to VPS-rich biofilm-associated cells, where productive infection becomes possible.

Peru-2 also differs fundamentally from T7-like phages in the organization of its internal core. In T7, the internal core proteins form a highly ordered structure that must unfold to pass through the narrow portal–tail lumen during infection. By contrast, Peru-2 contains a largely disordered internal core in mature particles. This intrinsic disorder may provide a functional advantage by allowing internal proteins to pass through the tail lumen without extensive unfolding, thereby facilitating rapid deployment and large-scale conformational rearrangements during infection initiation. Consistent with this model, we capture an open-tail intermediate in which a density corresponding to the P2_38 core protein lines the expanded tail lumen, supporting a deployment mechanism analogous to – but structurally more flexible than – that of T7 (42). The esterase lipase P2_42 is abundant in CsCl-purified virion preparations but is not resolved in cryo-EM structures. We speculate that this protein instead resides within the disordered internal core.

Importantly, the flagellar interactions observed for Peru-2 are mechanistically distinct from those of classical flagellotropic phages such as *Salmonella* phage χ (65, 66) and *Bacillus* phage PBS1 (67). In those systems, a specialized tail fiber binds the flagellar filament and harnesses its rotation to transport the phage directionally toward the cell body. By contrast, Peru-2 engages sheathed flagella, and genome ejection into this structure would likely be nonproductive. Instead, the sheathed flagellum may function as a dynamic scaffold that increases local virion concentration and promotes exploratory surface sampling prior to irreversible attachment elsewhere on the cell. Such a strategy may enhance adsorption efficiency by increasing the effective surface area available for phage attachment, particularly in motile cells where the flagellum extends far beyond the cell body. Consistent with this interpretation, comparisons between flagellated and aflagellated cells revealed only minor differences in phage titer in plate assays (data not shown).

Together, these structural and functional analyses show Peru-2 has evolved a distinct infection mechanism. Whereas T7-like and related ICP3 phages likely follow a similar and relatively linear infection pathway involving direct receptor binding followed by internal core protein ejection and genome release, Peru-2 employs a multimodal adsorption strategy (**Fig. 7**). Reversible flagellar binding may enhance encounter frequency; enzymatic tailspikes locally degrade the VPS barrier; and a conserved T7-like tail machine mediates final attachment and genome delivery upon interaction with a yet-unidentified outer membrane receptor. This hybrid architecture enables Peru-2 to retain the efficiency of the T7 infection core while incorporating accessory modules adapted to the structurally complex surface of *V. cholerae*. More broadly, Peru-2 illustrates how short-tailed T7-like phages can diversify through modular innovations, including Ig-like domains, biofilm-targeting tailspikes, and a disordered internal core, highlighting the remarkable evolutionary plasticity of phage infection strategies.

## Methods and Materials

### Bacteria and phage preparation

Bacteriophage Peru-2 was isolated during a cholera outbreak in 1993 in Peru and first characterized by Nicholas Judson in the John Mekalanos laboratory (30). *V. cholerae* strain C6706 was isolated by the CDC in 1991 in Peru. A spontaneous streptomycin resistant C6706 strain previously isolated was used to propagate Peru-2 in this work (68).

Peru-2 was grown by mixing phages with C6706 grown to late-log in a LB soft agar overlay on large LB agar plates (150 mm diameter). The LB media recipe used for strain growth is altered to be low salt (10 g Tryptone, 5 g yeast extract, and 5 g NaCl in distilled water). Phages were plated at a series of dilutions. Plates with confluent lysis resulting from dense and abundant small plaques were selected for phage preparation. 20-30 ml LB was first added to each plate and soft agar and liquid was scraped and collected. Both soft agar and liquid from 4-5 plates was collected and allowed to sit on ice for an hour for phage to diffuse out of soft agar. Liquid was then separated from agar and cells by centrifugation (5000× g, 10 minutes) and the supernatant filtered through a large 0.22 µM filter into a sterile bottle (Corning 321097). The supernatant was made 1.0 M NaCl and PEG 8000 was added to 8.0 % and dissolved by stirring at 4 °C. After 2-3 hours on ice, this supernatant was centrifuged at 8000× g for 15 minutes. The supernatant was discarded, and a diffuse pellet was suspended into 3.8 ml ice-cold TM buffer (Tris-HCl pH7.6 (10 mM) and MgCl_2_ (10 mM)). After incubating 60 minutes on ice, this was centrifuged at 5000× g for 10 minutes. The supernatant with dissolved phage was kept for CsCl density gradient purification.

Phage purification by CsCl density gradient centrifugation was performed in two steps. Initially, three step gradients of CsCl in TM buffer (ρ=1.43 (1.0 ml), 1.52 (0.5 ml), and 1.62 (0.5 ml)) were overlaid with 3.5 ml supernatant in 5.0 ml thin-walled Ultra-Clear tub (Beckman Coulter #344057). This was centrifuged at 40,000× rpm using a SW55.1 swinging bucker rotor for 90 minutes 4 °C. The phage band at ρ=1.42/1.62 interface was removed and added to 5.0 ml ice-cold CsCl in TM buffer (ρ=1.48) and mixed. This was centrifuged at 40,000× rpm for 16 hours. A single band was visible and removed for analysis.

### Phage genomics

The complete sequence and NCBI-annotated Peru-2 genome is deposited (NCBI accession number PX516383). We compared Peru-2 to both ICP3 isolate M1 (ON464735) and T7 (NC_001604) using EasyFig (v2.2.5) (69) and tblastx evalue cutoff of 0.05.

### Protein extraction for SDS-PAGE analysis and mass spectrometry proteomics

Proteins were extracted from phage particles using a method using methanol and chloroform (70). The protein pellet was dried and suspended in a mixture of water and actetonitrile (1:1), but did not completely dissolve. An aliquot was boiled in 1× NuPAGE LDS Sample Buffer (Invitrogen NP0007) and 1× NuPAGE Sample Reducing Agent (Invitrogen NP0009). This was run alongside a marker in precast 4-12 % NuPAGE Bis-Tris Mini Protein Gel (NP0321BOX) and stained with Coomassie Blue R-250 stain (BioRad #1610436).

Protein identification by peptide mass spectrometry was performed by trypsin digestion of the sample and running peptides on a Thermo Orbitrap Exploris 480 coupled to a ThermoEASY-nLC1200 nano-UPLC at the Beth Israel Deaconess Medical Center-Harvard Spectrometry Facility.

### Phage titer and adsorption assays

Phages were serially diluted and titered on C6706 and two isogenic mutants to measure the concentration of infectious particles using LB agar and a 3.5mL soft LB agar overlay (0.5%) in 100mm petri dishes. In brief, 200 μL of a bacterial culture in late-log growth at 37°C was mixed with 100 μL of diluted phage prior to plating. For phage adsorption assays, 100 μL of 1.0 × 10^10^ PFU/mL phage stock was mixed with a 5.0 mL bacterial culture at OD 1.0 (MOI ∼1.0) and the infected culture was grown shaking at 37°C. At defined time points, a 1.0ml aliquot was removed, quickly centrifuged at 15,000 RPM and the supernatant was titered as above on C6706.

### Cryo-EM sample preparation and data collection

The ultracentrifuged Peru-2 phages with CsCl were dialyzed 20 mM Tris pH 7.6, 500 mM NaCl at 4 °C overnight. 3 µL dialyzed phages at OD_260_ of 1.0 were placed onto predischarge Quantifoil R2/1 Cu300 mesh grid then vitrified in liquid ethane blotted 5 seconds using GP2 (Leica) stored at liquid nitrogen until loaded into microscope.

The Peru-2 phages were imaged using a Titan Krios G2 electron microscope (Thermo Fisher Scientific) equipped with a field emission gun, a K3 summit direct electron detector (Gatan), and an energy filter with a slit width of 20 eV. SerialEM (71) was used to collect cryo-EM image stacks with a defocus range of −1.0 to −2.0 µm, receiving a total dose of ∼55 e^−^/Å^2^ distributed over 40 frames (0.07 s/frame). With a nominal magnification of 81,000×, 6,621 movies were collected in total with a pixel size of 0.534 Å at super resolution mode.

### Cryo-EM single-particle analysis

Flowcharts of the cryo-EM data-processing pipeline are shown in ***SI Appendix*, Fig. S1**. Dose-fractionated movies were initially processed in cryoSPARC (72) by patch motion correction and patch CTF estimation, with images cropped to a calibrated pixel size of 1.068 Å. A total of 484 manually picked phage capsid particles were used to generate templates for Topaz training (73). Following automated particle picking with Topaz, particle extraction, and 2D classification, 11,395 empty capsid particles and 187,061 DNA-filled capsid particles were selected for further analysis. Selected particles were imported into RELION (74), and corresponding micrographs were reprocessed using CTFFIND4.1 for CTF estimation (75). Particles were re-extracted at bin 4 and subjected to 3D refinement with I3 symmetry imposed, followed by focused 3D classification on a single capsid vertex. This procedure yielded 3,789 tail-containing particles from empty capsids and 130,625 tail-containing particles from DNA-filled capsids. For both datasets, the capsid vertex opposite the tail was extracted and locally refined with C5 symmetry, resolving the capsomer and associated minor capsid proteins to 2.35 Å resolution using 2,079,915 particles.

To analyze the tail complex, particles were subjected to C5 symmetry expansion and 3D classification in cryoSPARC (72) to overcome the intrinsic C5 symmetry imposed by the capsid vertex. Tail particles from DNA-filled capsids separated into two distinct conformational states. A closed state was refined to 3.18 Å from 33,052 particles, and an open state was refined to 3.61 Å from 25,567 particles. Both reconstructions were obtained through iterative rounds of local CTF refinement and local refinement with C6 symmetry. In contrast, tail particles from empty capsids yielded a 3.92 Å reconstruction from 2,530 particles after local refinement with C6 symmetry, in which density corresponding to the portal barrel was restored.

Finally, C6 symmetry expansion was applied to both tail conformations to extract side tail fibers. Heterogeneous conformations were identified by 3D classification, and the most populated class was refined to 3.65 Å resolution from 321,124 particles using local refinement.

### Model building and refinement

AlphaFold3 (76) was used to generate initial atomic models of the proteins based on the amino acid sequence. The models were fitted with map densities enhanced by EMready (77) and automatically refined using Namdinator (78). Coot (79) was used to perform manual refinements on the models. phenix.real_space refine (80) was also used to perform final model refinements. Model refinement, and validation are listed in ***SI Appendix*, Table S1**.

### Cryo-ET sample preparation, data collection, and reconstruction

*V. cholerae* wild-type and Δ*flhG* strains were grown in LB medium to an OD_600_ of 1.0. For infection, 20 µL of each culture was mixed with 3 µL of Peru-2 phage and incubated for 30 min at 37 °C. Aliquots (5 µL) of the mixtures were applied to Quantifoil R2/1 Cu300 mesh grids and vitrified in liquid ethane using a GP2 (Leica), with an 8-second blot time. Tilt series of bacteria infected by phages were collected using SerialEM (71) on a Titan Krios G2 microscope (Thermo Fisher Scientific) equipped with a field-emission gun, a K3 Summit direct electron detector and an energy filter with a slit width of 20 eV (Gatan). Data were acquired using FastTomo (81) at a nominal magnification of 42,000×, corresponding to a calibrated pixel size of 2.148 Å, with a defocus of approximately 5 µm. Tilt series were collected from −48° to +48° with a 3° increment. Tilt series alignment and tomograms reconstruction were performed using IMOD (82). In total, 88 tomograms were generated for WT cells and 50 tomograms for the Δ*flhG* mutant.

## Data availability

The electron density maps, and atomic coordinates have been deposited in the Electron Microscopy Data Bank and Protein Data Bank under accession codes: EMD-74388 (major capsid and minor capsid), EMD-74389 (closed tail complex), EMD-74616 (open tail complex), EMD-77075 (tailspike), and the PDB code PDB-9ZKX (major capsid and minor capsid), PDB-9ZKY (closed tail complex), PDB-9ZRH (open tail complex), PDB-13HW (tailspike).

## Acknowledgments

We thank Jennifer Aronson for critical reading of the manuscript. We also thank Wangbiao (Seven) Guo for guidance on *V. cholerae* sample preparation and Chunyan Wang for cryo-EM sample screening. We acknowledge preliminary work on Peru-2 by Nicholas Judson (John Mekalanos Laboratory at HMS) and Aaron Gray (Ian Molineux Laboratory at UT Austin) which helped in developing methods for Peru-2 growth and purification. We thank John Asara at the BIDMC-Harvard mass spectrometry facility for his assistance in identifying proteins by mass spectrometry. This work is supported by NIH grants R01GM124378, 5R37AI018045, and R01GM110243; cryo-EM data were collected at Yale Cryo-EM Resources funded in part by NIH grant 1S10OD023603-01A1.

