## Supplemental Figures for "Structural basis of multimodal adsorption and infection initiation by *Vibrio* phage Peru-2"

### Supplemental Information

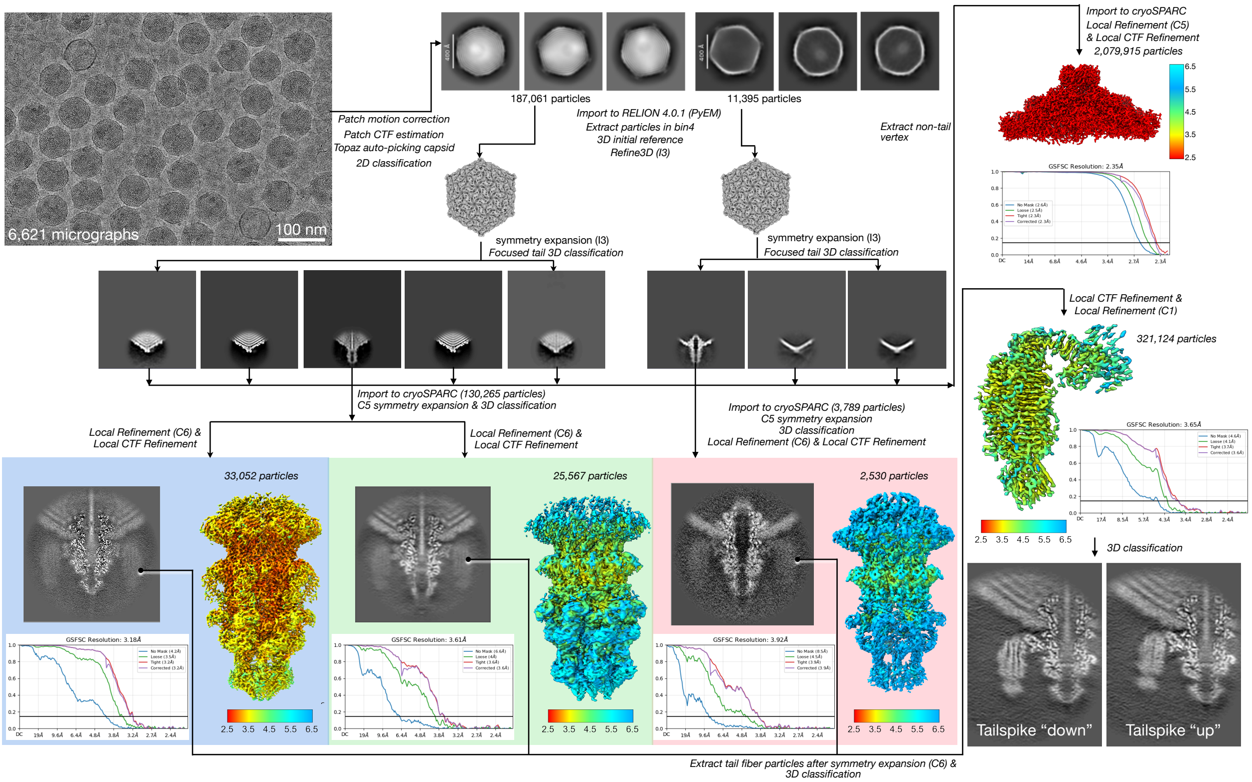


***SI Appendix*, Fig. S1 | Cryo-EM data processing workflow from raw micrographs to near-atomic structure determination.** Tail orientations were determined using RELION 3D classification. Three distinct tail structures were subsequently reconstructed in cryoSPARC, with final resolutions estimated at the FSC 0.143 criterion. Local resolution estimation for each reconstruction are displayed as color-gradient maps.

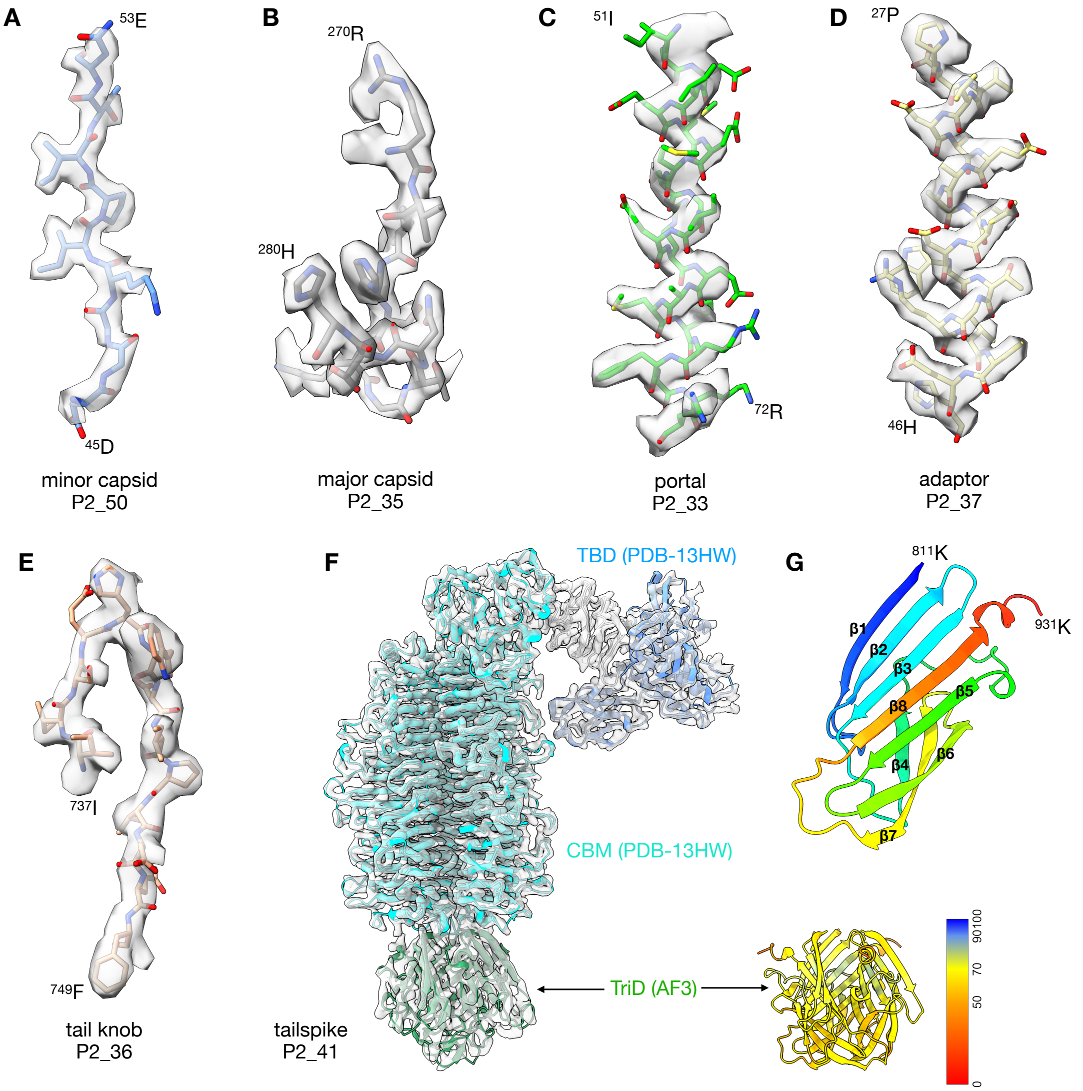


***SI Appendix*, Fig. S2 | Superimposition of the atomic models fitting into cryo-EM densities of pre-ejected mature phage with closed tail. (A-E)** Representative residues are shown for minor capsid, major capsid, portal, adaptor, and tail knob to illustrate good model-to-map fitting at near-atomic resolution. (**F**) *Left*: The tailspike shows an overall good fit to the cryo-EM density (EMD-77075). The secondary structure of TriD could not be built due to its flexibility and is therefore shown as an AlphaFold3-predicted model fitted into the map. *Right*: The AlphaFold3 prediction of the TriD is shown on the right, colored by model confidence. (**G**) The TriD monomer consists of eight β-strands arranged into two opposing β-sheets, containing five and three strands, respectively, that form a β-sandwich.

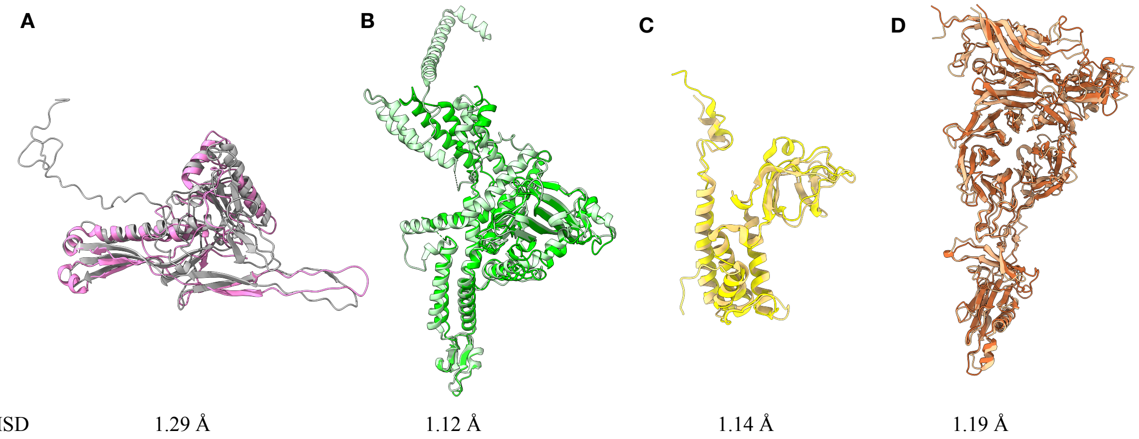


***SI Appendix*, Fig. S3 | Superposition of the structural models of mature Peru-2 and T7. (A)** Major capsid of Peru-2 (P2_35 in grey) and T7 (gp10 in pink, PDB-2XVR). **(B)** Portal of Peru-2 (P2_33 in green) and T7 (Gp8 in light green, PDB-9JYZ). **(C)** Head-tail adaptor of Peru-2 (P2_37 in yellow) and T7 (Gp11 in light yellow, PDB-9JYZ). **(D)** Tail knob of Peru-2 (P2_36 in brown) and T7 (Gp12 in orange, PDB-9JYZ). RMSD values are shown for the corresponding structural alignments between Peru-2 and T7 proteins.

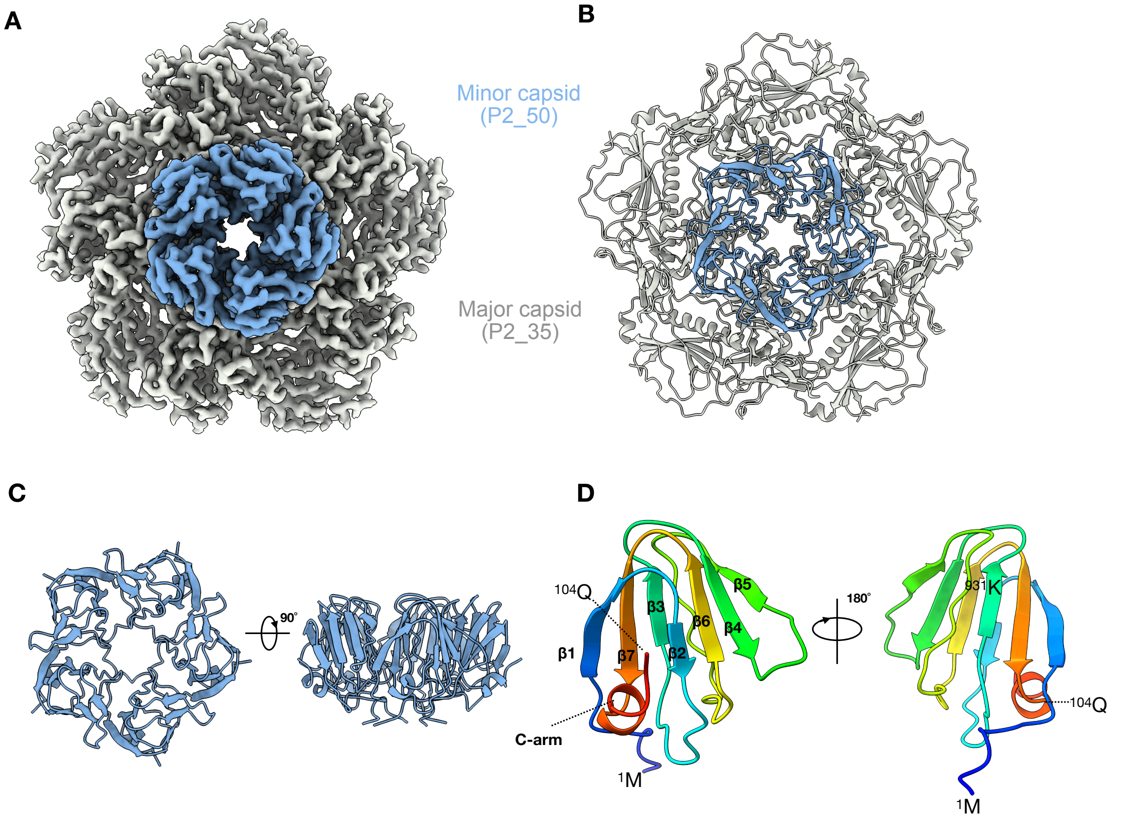


***SI Appendix*, Fig. S4 | Minor capsid protein p2_50 adopts an Ig-like domain.** (**A,B**) Top view of the P2_50 pentamer and Gp_35 pentamer cryo-EM density and ribbon model. (**C**) Top and side views of the Gp_35 pentamers. (**D**) The P2_50 monomer consists of seven β-strands and a C-terminal α-helical arm, together forming a classic Ig-like fold.

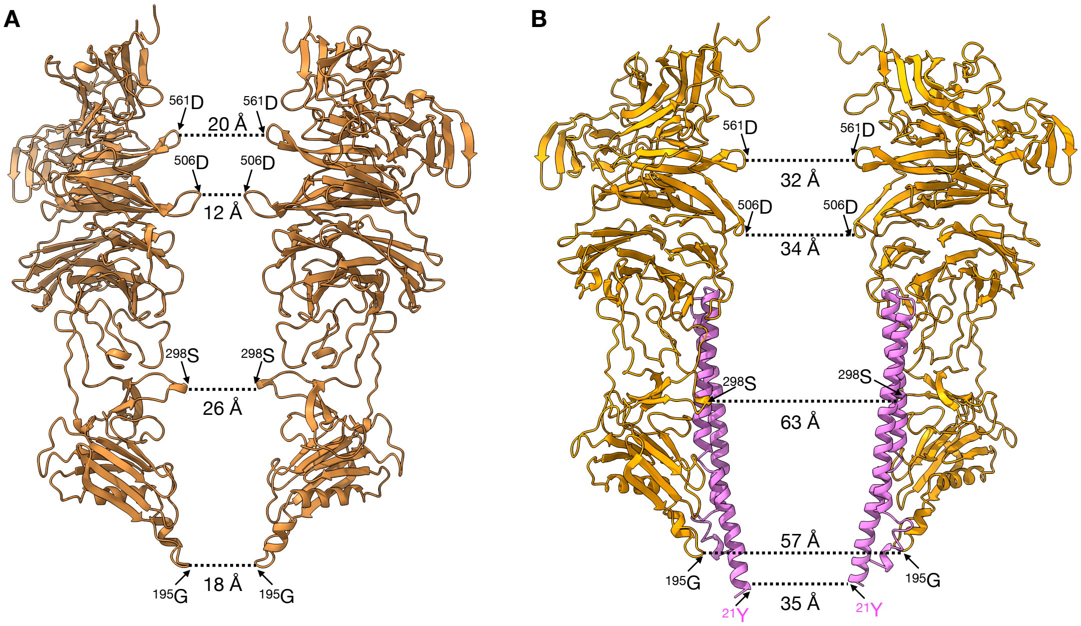


***SI Appendix*, Fig. S5 | Constriction-gate distances within the Peru-2 tail in the pre-ejection and ejection states. (A)** Distances measured at key constriction points (residues $^{195}$G, $^{298}$S, $^{506}$D, and $^{561}$D of the tail knob protein P2_36) in the pre-ejection state. **(B)** Corresponding measurements at the same constriction gates in the ejection state, illustrating the widening of the tail lumen to enable genome translocation. An internal core protein P2_38 is also shown, with the measurement taken at residue $^{21}$Y.

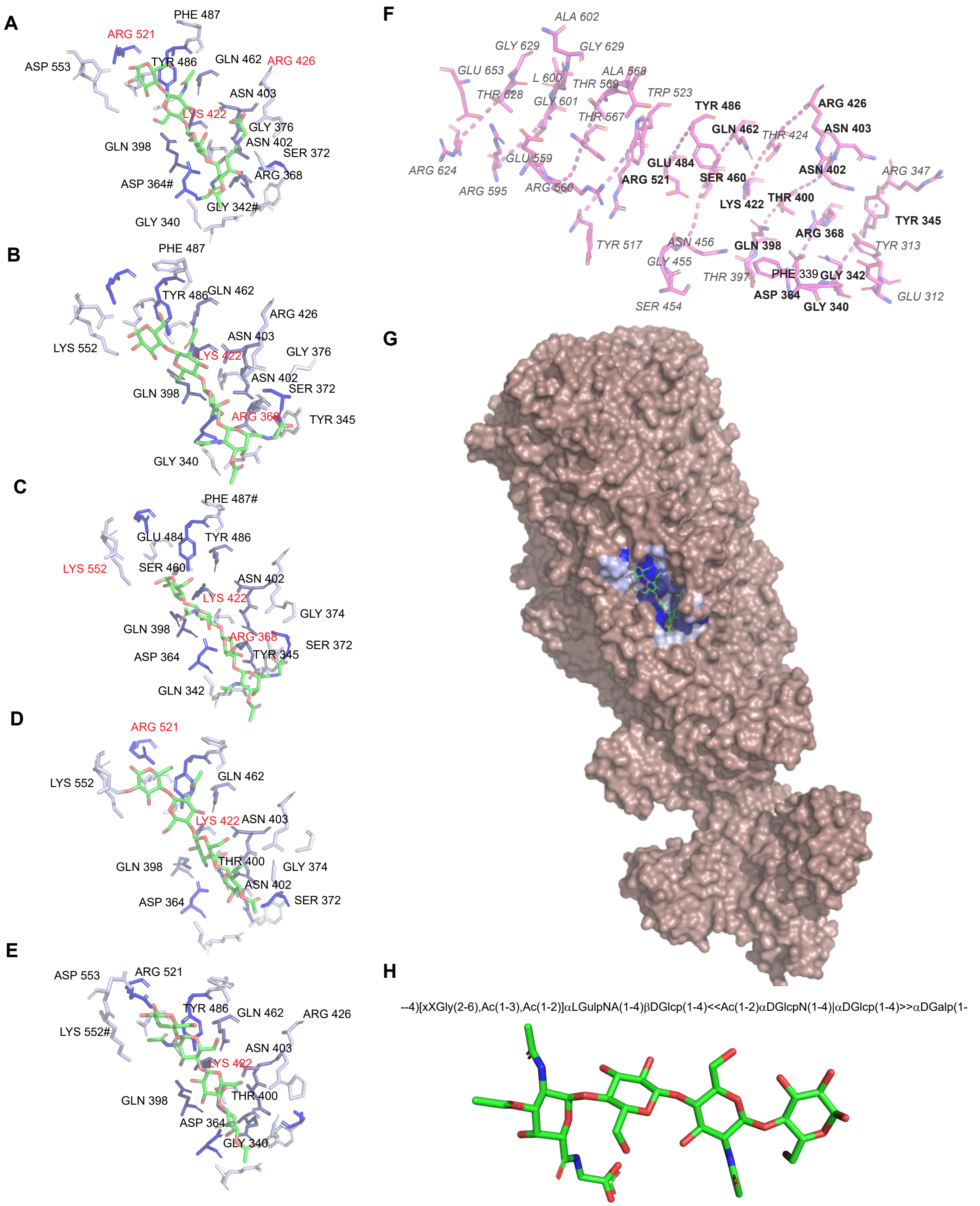


***SI Appendix*, Fig. S6 | *In silico* prediction of VPS binding in tailspike reveals potential molecular contacts.** (**A-E**) Protenix-based orientation of VPS unit into coordinate (PDB: 13HW) built by cryo-EM density. 5 distinct models were generated and VPS-interacting amino acids are labeled for each. Red-colored labels indicate predicted salt bridges and # indicates hydrophobic interactions. Other labels indicate Hydrogen-bond interactions. (**F**) GlycanInsight prediction of

VPS-binding pocket in Tail Spike. Residues that interact are labeled and shown. Residues that overlap with Protenex model are in bold and not italicized. (**G**) Cryo-EM structure of tailspike trimer with pocket residues colored in blue. Each residue is colored by the frequency of residue identification in the 5 distinct models. Light blue residues are sparsely represented the darkest blue are identified in all models. (**H**) IUPAC Condensed Nomenclature of the major polysaccharide unit in VPS and stick molecular representation.

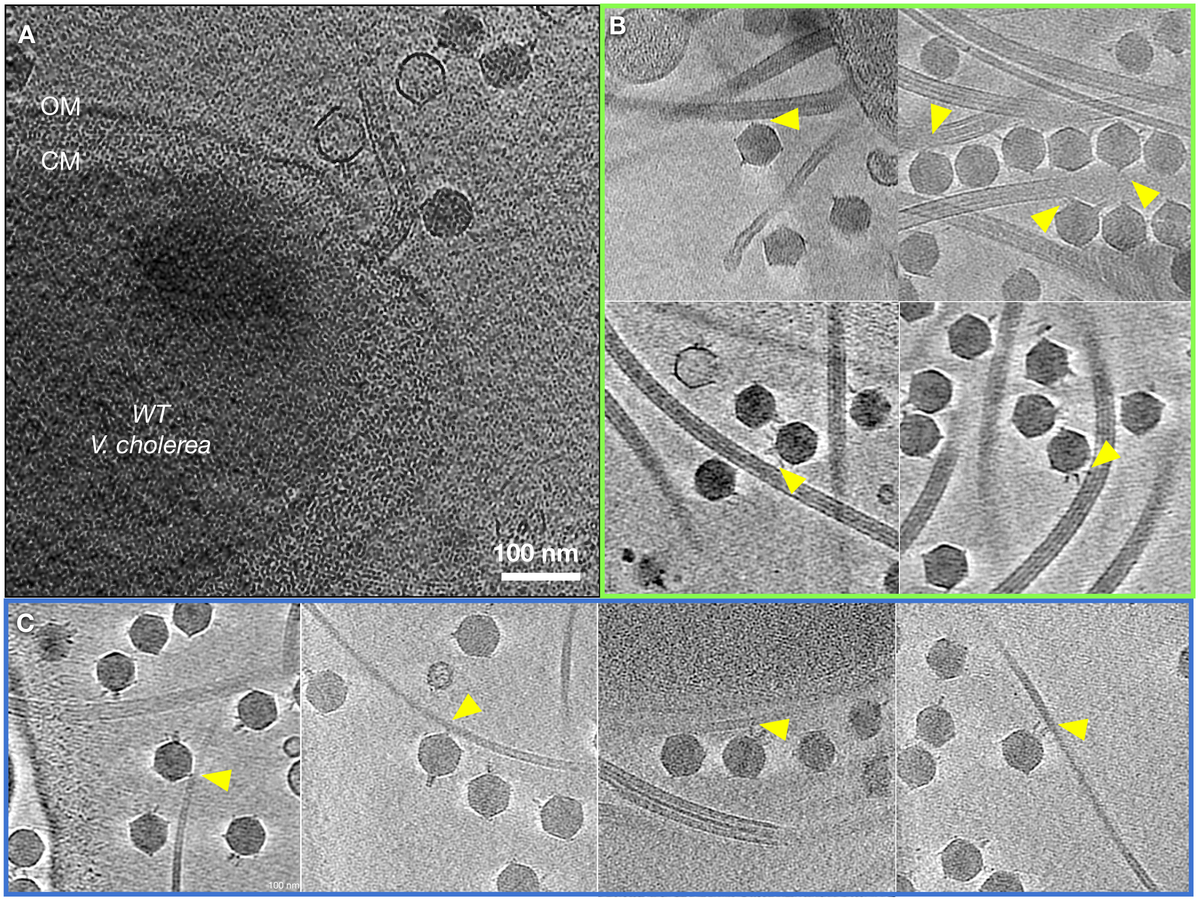


***SI Appendix*, Fig. S7 | Cryo-ET reveals multiple modes of Peru-2 adsorption. (A)** Representative tomograms of Peru-2 particles interacting with *V. cholerae* wild type. **(B)** Examples of initial adsorption events in which capsid and tailspikes engage the sheathed flagellum. **(C)** Initial adsorption events mediated by capsid and tailspikes on non-sheathed flagella. Yellow arrows indicate interaction sites or phage particles undergoing adsorption.

***SI Appendix*, Table S1 | Cryo-EM data collection, processing, model refinement, and validation.**

| | **capsid** | **closed tail** | **open tail** | **tailspike** |
|---|---|---|---|---|
| **Data collection** | | | | |
| Magnification | 81,000× | | | |
| Voltage (kV) | 300 | | | |
| Electron ($e^-$/Å$^2$) | 50 | | | |
| Defocus range (μm) | -0.6 to -1.2 | | | |
| Pixel size (Å) | 1.068 | | | |
| Micrographs (no.) | 6,621 | | | |
| **Data processing** | | | | |
| Symmetry imposed | C5 | C6 | C6 | C1 |
| Final particle images (no. ) | 2,079,915 | 33,052 | 4,452 | 321,124 |
| Map resolution (Å) 0.143 FSC | 2.35 | 2.87 | 3.92 | 3.65 |
| Map sharpening B factor (Å$^2$) | -101.1 | -61.0 | -16.7 | -105.9 |
| **Refinement** | | | | |
| CC (model vs. data) | 0.75 | 0.81 | 0.71 | 0.80 |
| Chain count | 10 | 30 | 12 | 3 |
| Non-hydrogen atoms | 16,580 | 100,074 | 43,608 | 18,991 |
| Protein residues | 2,170 | 12,594 | 5,550 | 2,430 |
| Ligands | 0 | 0 | 0 | 0 |
| B factors (Å$^2$) | | | | |
| Proteins | 22.91 | 85.95 | 115.24 | 113.60 |
| Ligands | / | / | / | / |
| R.m.s. deviations | | | | |
| Bond lengths (Å) | 0.004 (0) | 0.003 (0) | 0.004 (0) | 0.024 (17) |
| Bond angles (˚) | 0.543 (1) | 0.711 (49) | 0.680 (17) | 0.729 (14) |
| Validation | | | | |
| MolProbity score | 1.32 | 1.93 | 2.24 | 2.19 |
| Clash score | 4.11 | 7.54 | 5.57 | 6.85 |
| Ramachandran plot | | | | |
| Favoured (%) | 97.71 | 96.07 | 93.67 | 92.99 |
| Allowed (%) | 2.29 | 3.77 | 6.19 | 6.85 |
| Outliers (%) | 0.00 | 0.16 | 0.14 | 0.17 |
| PDB code | 9ZKX | 9ZKY | 9ZRH | 13HW |
| EMDB code | EMD-74388 | EMD-74389 | EMD-74616 | EMD-77075 |
